# Sex differences in reward- and punishment-guided actions

**DOI:** 10.1101/581546

**Authors:** Tara G. Chowdhury, Kathryn G. Wallin-Miller, Alice A. Rear, Junchol Park, Vanessa Diaz, Nicholas W. Simon, Bita Moghaddam

**Affiliations:** Oregon Health and Science University Department of Behavioral Neuroscience; University of Pittsburgh Department of Neuroscience

## Abstract

Differences in the prevalence and presentation of psychiatric illnesses in men and women suggest that neurobiological sex differences confer vulnerability or resilience in these disorders. Rodent behavioral models are critical for understanding the mechanisms of these differences. Reward processing and punishment avoidance are fundamental dimensions of the symptoms of psychiatric disorders. Here we explored sex differences along these dimensions using multiple and distinct behavioral paradigms. We found no sex difference in reward-guided associative learning but a faster punishment-avoidance learning in females. After learning, females were more sensitive than males to probabilistic punishment but less sensitive when punishment could be avoided with certainty. No sex differences were found in reward-guided cognitive flexibility. Thus, sex differences in goal-directed behaviors emerged selectively when there was an aversive context. These differences were critically sensitive to whether the punishment was certain or unpredictable. Our findings with these new paradigms provide conceptual and practical tools for investigating brain mechanisms that account for sex differences in susceptibility to anxiety and impulsivity. They may also provide insight for understanding the evolution of sex-specific optimal behavioral strategies in dynamic environments.

## INTRODUCTION

Men and women show different rates of diagnosis, symptomology, and treatment responsivity in most brain disorders. For instance, men are more commonly diagnosed with schizophrenia, four times more likely to suffer from attention-deficit/hyperactivity disorder (ADHD) (Rowland, Lesesne, & Abramowitz, 2002), and twice as likely to be currently abusing illicit drugs (Abuse, 2013). On the other hand, major depressive disorder (MDD) and anxiety disorders are more common in women (Cyranowski, Frank, Young, & Shear, 2000). These patterns suggest that biological sex differences may underlie the disparities in vulnerability to these illnesses.

Reward-seeking and punishment-avoidance behaviors are fundamental to motivation, and are critical components of the pathophysiology of most psychiatric illnesses. For example, studies have shown that patients with depressive disorders (Pizzagalli et al., 2009), schizophrenia (Juckel, 2016; Kirsch, Ronshausen, Mier, & Gallhofer, 2007; Schlagenhauf et al., 2008), as well as ADHD (Scheres, Milham, Knutson, & Castellanos, 2007; Strohle et al., 2008) have modified brain responses to reward. In particular, major depression is associated with both a hyposensitivity to rewarding stimuli and hypersensitivity to punishment, and associated imaging findings are correlated with increasing frequency of depressive episodes (Kumar et al., 2018). Importantly, disorders where reward and aversive responding are differently affected also show different incidences and symptom profiles in women versus men. For example, in anxiety disorders there is an increased avoidance of aversive stimuli that is more pronounced in women (Sheynin et al., 2014). Similarly, in MDD, men are more likely to exhibit symptoms of aggression, substance abuse, and risky behavior, while females report higher rates of sleep disturbance, stress, and anhedonia (L. A. Martin, Neighbors, & Griffith, 2013). Similarly, it appears that sex influences substance abuse patterns, with women more likely than men to cite stress as the reason for initiating or relapsing drug use (Becker, McClellan, & Reed, 2017).

Finally, in ADHD, females diagnosed with the disorder more frequently experience comorbid anxiety and depression. These findings suggest that distinct responses to punishment may explain some of the observed sex differences in both the prevalence and expression of mental disorders.

Animal models are critical for understanding the behavioral neuroscience of sex differences, allowing assessment of motivated behaviors relevant to psychiatric disorders. Animal behavioral models, however, have been primarily developed and characterized with only male subjects. Additionally, these models often assess reward and punishment contingencies in separate tasks and constructs. In naturalistic environments, appetitive and aversive contexts and outcomes are often intertwined, and the ability to accurately assess and respond to these conflicting situations is critical for optimal decision-making. Females have been shown to respond differently than males to punishing stimuli (Denti & Epstein, 1972; Gruene, Flick, Stefano, Shea, & Shansky, 2015; Orsini, Willis, Gilbert, Bizon, & Setlow, 2016; Voulo & Parsons, 2017) as well as stress (Bangasser & Valentino, 2014; McEwen, 2014); however, little is known about how sex differences translate to tasks that involve both reward and punishment in the same behavioral series. Here, we investigate sex differences in multiple rodent behavioral tasks involving motivated behaviors (defined as behaviors where actions are guided by action-outcome contingencies) that integrate reward-seeking and punishment avoidance.

In the first task, the Punishment Risk Task (PRT), a rewarded action was associated with an escalating probability of punishment (Park & Moghaddam, 2017). Thus, action-reward contingency was certain but different blocks in the same behavioral series were associated with varying probability of receiving a shock after action execution. This task assesses behavior in response to unpredictability and perception of potential threats, which is relevant to human models of anxiety (Cornwell, Garrido, Overstreet, Pine, & Grillon, 2017). Human sex differences in prevalence and expression of anxiety-related behaviors suggest that males and females may respond differently to anxiety-provoking situations. This task, therefore, allowed us to compare motivated actions in male and female rats within a context of anxiety.

A second novel task, the Approach Avoid Task (AAT) measured actions to seek reward or avoid punishment during the same behavioral session. Rats were given simultaneous access to two distinct actions (lever-press or nose-poke), and two discriminative stimuli (a light or tone cue) signaled the trial type, either approach, in which a specific action was reinforced with a pellet, or avoidance, in which the other action prevented onset of a foot-shock. This task is relevant to symptoms of brain disorders such as anxiety, substance abuse, and MDD, which are associated with differences in processing of rewarding and punishing contexts (Dombrovski, Szanto, Clark, Reynolds, & Siegle, 2013; McCabe, Woffindale, Harmer, & Cowen, 2012). In addition to comparing learning and performance of male and female rats in this task, we conducted two experiments to gauge the impact of traditional models of anxiety in the context of this paradigm. These included measuring (1) the dose-response effect of a pharmacological model of anxiety (anxiogenic drug FG7142) on performance of the AAT and (2) elevated plus maze (EPM) performance of animals that had undergone training on the AAT compared to naïve animals.

Finally, we characterized sex differences in reward-motivated tasks that required behavioral inhibition and flexibility. Most disorders with higher prevalence in men than women (autism, ADHD, schizophrenia) involve impaired behavioral inhibition and flexibility. In addition, male predisposition to compulsive behavior may contribute to the higher rates of substance abuse and dependence in men [20]. To investigate these constructs, we used two operant tests of cognitive flexibility: reversal learning and extradimensional shifting. Both tasks required subjects to update behavior in response to changing rules.

## METHODS

All of the animal procedures were performed in accordance with the National Institutes of Health Guide for the Care and Use of Laboratory Animals and approved by the Animal Welfare Committee at the University of Pittsburgh and Oregon Health and Science University. Separate cohorts of animals were used for each task.

### Task 1: Punishment Risk Task (PRT)

#### Animals

Six female and 6 male Long Evans rats were delivered to the Oregon Health and Science University animal facility at the age of 57 days. Animals were housed with *ad libitum* food and water in temperature-controlled and humidity-controlled rooms on a 12 h on/12 h off reverse light/dark cycle with lights on at 1900 hours. After 1 week of acclimatization, animals were handled and weighed daily. Animals were food restricted to 90% of their free-feeding body weight. Animals were fed once daily after completion of the behavioral tasks.

#### Punishment Risk Task (PRT)

This task followed methods previously described in (Park & Moghaddam, 2017). Briefly, an operant chamber was equipped with a nose-poke port on one side and a food trough on the opposite side (Figure 1A). Rats were trained to make an instrumental response at the nose poke port to receive a sugar pellet at the food trough on the fixed ratio schedule of one (FR1). After successful completion of three FR1 training sessions consisting of 90 trials in 60 mins, rats underwent a no-shock session in which they performed 90 trials divided into three blocks separated by two minute time outs in the absence of punishment. Task 1 sessions consisted of three blocks of 30 trials each. Rats were trained for 2 days on the behavioral task prior to the recording session. Throughout the sessions, action-reward contingency remained constant, with one nose-poke resulting in one sugar pellet. In contrast, the probability of receiving a punishment paired with the reward increased over the blocks (0, 6 or 10% in blocks 1, 2 and 3, respectively). Punishment was a brief foot shock (300 ms electrical shock of 0.3 mA).

**Figure 1:**
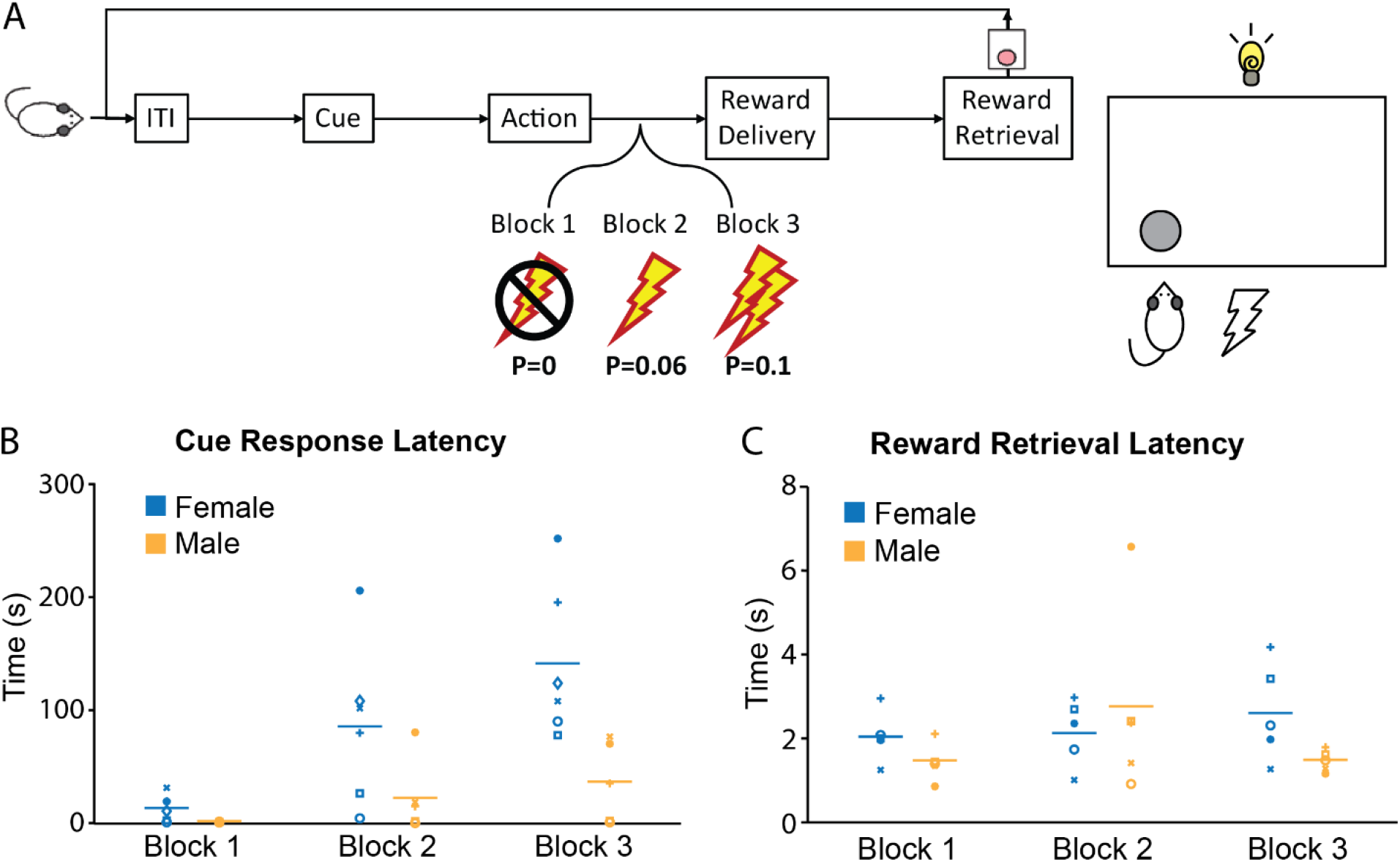
Sex differences measured during reward-seeking with a variable risk of punishment in the Punishment Risk Task. Reward seeking behavior was measured in a FR1 task in three blocks of 30 trials. The first block was a standard FR1 task with no risk of punishment, followed by two blocks that introduced risk of a foot shock immediately after the rewarded action and before reward delivery (A). In females, block 1 response times were significantly shorter than block 2 (p<0.0038) as well as block 3 (p<0.001), with an additional decrease between blocks 2 and 3 (p=0.0349). There were no significant differences in latency between any of the individual blocks in males. When male and female groups were combined, we found a significant effect of block (B; RM-ANOVA: F (2, 18) = 15.8; *p* = 0.0001), with increasing cue response latency as the shock probability increased. Significant differences were found between blocks 1 and 2 (p=0.0075), 2 and 3 (p=0.0611), and 1 and 3 (p<0.0001). RM-ANOVA indicates a significant effect of sex (F(1,9)=6.949;P=0.0271) with an additional interaction of sex and block (F(2,18)=5.044; p=0.0182). No significant differences were found in reward retrieval latency (C).

To minimize generalization of the action-punishment contingency, blocks were organized in an ascending shock probability with 2-min timeouts between blocks. Punishment trials were pseudo-randomly assigned, with the first shock occurring within the first 5 trials. All training sessions were terminated if not completed in 180 mins. Animals displayed stable behavioral performance without signs of contextual fear conditioning, as they performed without signs of anxiety in block 1 (with 0% punishment) across all sessions. Figure 1A illustrates operant chamber set up. One side was equipped with a sugar pellet dispenser and food trough with infrared (IR) beams to detect retrieval of pellets. The opposite side was fitted with a closing nose-poke port and a retractable lever. Additionally, the chamber was equipped with an audible tone module and a salient cue-light was positioned on the wall with the nose-poke and lever. The floor of the box was an electrified grating for delivery of foot-shocks.

All behavioral sessions were conducted between 1200 and 1900 hours. Animals were handled in the behavioral testing room and given sugar pellets for two days prior to the beginning of training.

#### Exclusion of data points

Two females failed to complete the entire session within the allotted 180 min. For the results reported, data from all *completed* trials were averaged and included in analysis; therefore, the time between the last completed nose-poke and the end of the session was excluded. When these individuals were excluded from the analysis entirely (N=5M, 4F), the effect of shock on response latency remained significant (*p* = 0.0002), while the effect of sex (*p* = 0.06) and the interaction between sex and shock (*p* = 0.086) trended toward significance.

### Task 2: Approach Avoid Task (AAT)

#### Animals

Eleven male and 11 female Sprague-Dawley rats were delivered to the University of Pittsburgh animal facility at the age of 57 days. Animals were housed with *ad libitum* food and water in temperature-controlled and humidity-controlled rooms on a 12 h on/12 h off reverse light/dark cycle with lights on at 1900 hours. After 1 week of acclimatization, animals were handled and weighed daily. Animals were food restricted to 90% of their free-feeding body weight. Animals were fed once daily within 2 hours of the end of the dark cycle.

Six male and 6 female Sprague-Dawley rats were delivered to the Oregon Health and Science University animal facility at the age of 57 days. Animals were housed with *ad libitum* food and water in temperature-controlled and humidity-controlled rooms on a 12 h on/12 h off reverse light/dark cycle with lights on at 1900 hours. After 15 days of acclimatization, these animals were tested on the EPM, as described below.

#### Magazine Training

Day 1 consisted of a 30-min magazine training session. Each trial began with the delivery of a sugar pellet to the food trough and ended when the animal broke the IR beam in the food trough by retrieving the pellet. The inter-trial interval (ITI) was variable, with an average of 45 s.

#### Action for Reward

Rats were trained to perform a single nose-poke to obtain a sugar pellet reward. Each trial began with the nose-poke port open and illuminated. When the animal performed a nose-poke, a sugar pellet was immediately delivered into the food trough, and the nose-poke port was closed. The trial concluded with retrieval of the sugar pellet at the food trough. The ITI was variable, with an average of 17 s. Criterion was 50 trials completed within 90 minutes, after which rats moved on to ‘lever-press for reward’ or ‘nose-poke to avoid’ training. Lever-press for reward training was identical to nose-poke for reward with the exception that the nose-poke port stayed closed and the retractable lever was made available at the beginning of each trial and retracted upon a successful lever-press.

#### Action to Avoid

Action to avoid training consisted of hand-shaping the animals to nose-poke or press the lever to stop a foot-shock. During this training session, the nose-poke or lever remained in the available position, and a hand-shaping switch was connected to control the foot-shock grating. When the switch was pressed, a 0.3 mA shock was delivered to the grating. The animal was guided to perform the correct action in a stepwise fashion, with the experimenter stopping the shock when the animal was on the correct side of the box, then the correct quadrant, then when the animal faced the nose-poke or lever, and finally when the animal’s nose was near the nose-poke or paw was on the lever. The training was considered complete when the animal successfully stopped the shock by performing the correct action for 5 instances of the shock in a row. Following successful action to avoid training, the animals went on to cued action for reward training.

#### Cued-Action-for-Reward

Cued-action-for-reward trials began with a 10-s cue, either a tone or a light, during which the nose-poke or lever was made available. After 10 s, the cue would turn off and the nose-poke port would close or the lever would retract. If the animal performed a successful action during the cue, a sugar pellet reward was delivered, and the trial would end upon retrieval of the reward. Each session consisted of 50 trials with a variable ITI of average 17 s. Animals continued daily action for reward sessions until they reached a criterion of 70% successful trials (at least 35 out of 50 trials). Once animals reached criterion, they went on to cued action to avoid training, or the full task if both training types were complete.

#### Cued-Action-to-Avoid

Cued-action-to-avoid trials began with a 10-s cue, either a tone or a light, during which the nose-poke or lever was made available. If the animal performed the action during the 10-s cue, the cue would end and the nose-poke port would close or the lever would retract, and the trial would be considered a successful “avoid”. If the animal did not perform the action before the 10-s cue ended, the foot-shock would begin and continue for 10 s or until the animal performed the action, considered an “escape” response. Each session consisted of 50 trials with a variable ITI of average 17 s. Animals continued daily action to avoid sessions until they reached a criterion of 70% successful trials (at least 35 out of 50 trials).

#### Approach Avoid Task (AAT)

Once animals successfully reached criterion on both cued approach and avoid training types, they began training in the full AAT (Figure 2A). During a session, the two trial types were interleaved in a pseudorandom fashion until 50 trials of each type had been completed. At the onset of the cue, both the nose-poke and lever were made available, if the animal performed the incorrect action during a reward trial, the trial ended with no reward delivered. If the animal performed the incorrect action during an avoid trial, the foot-shock would immediately begin, but animal was still allowed to escape the shock with the correct action. Animals continued training until their behavior was stable for at least 5 days.

**Figure 2:**
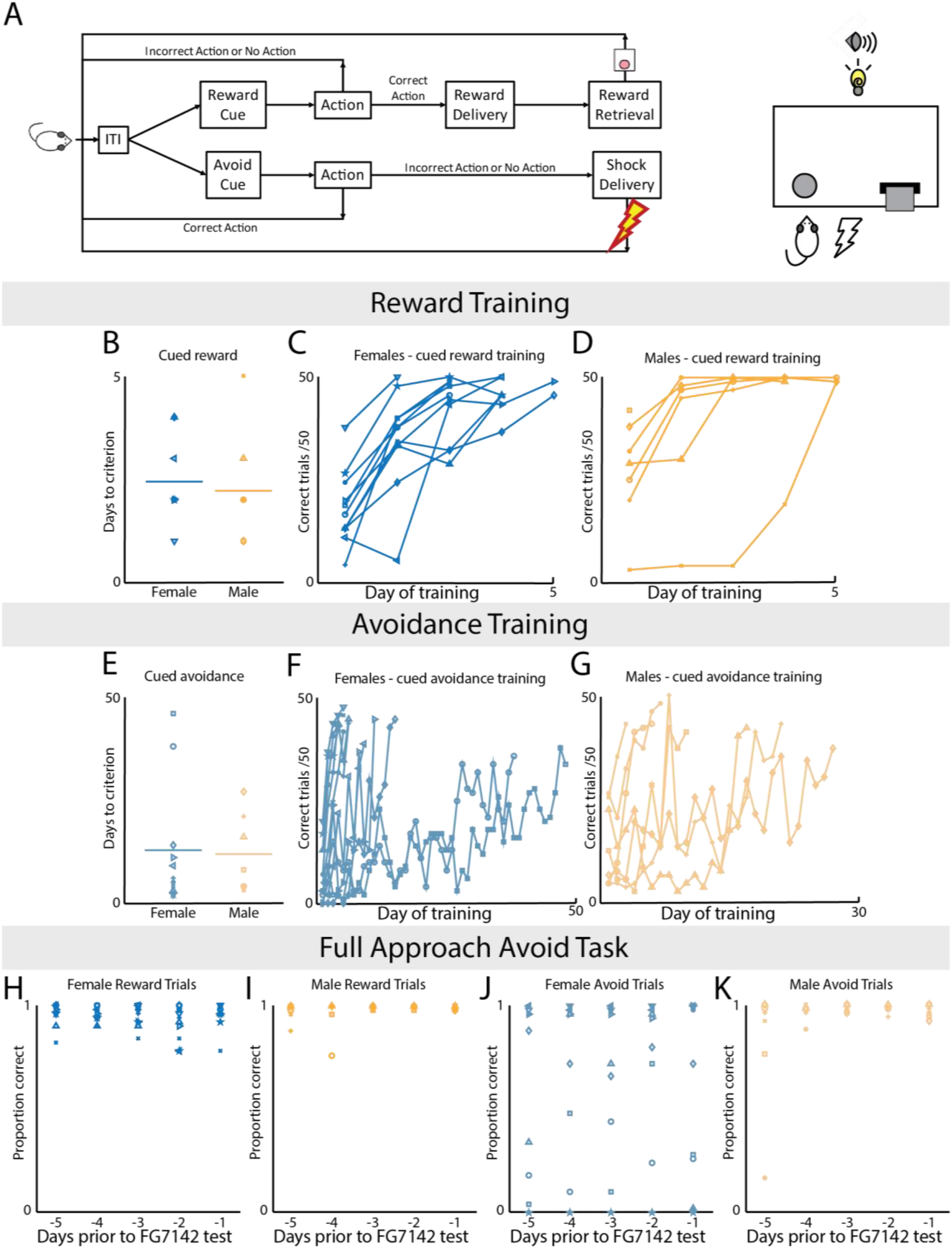
Sex differences in acquisition of reward seeking and punishment avoidance behaviors. Rats were initially shaped on the reward-seeking and punishment-avoidance components of the task separately, before being trained on the combined full task (A) until stable performance emerged. Cued-action-for-reward performance was observed daily until animals achieved a criterion of 70% correct trials (B shows days to criterion of all individuals). Average and individual performance in females (C) and males (D) is shown, and sex differences were not observed in reward learning acquisition (males: 2.1 ± 0.4 days; females: 2.5 ± 0.3 days; t(16) = −0.583; *p* = 0.568). Cued-action-to-avoid performance is shown in similar fashion (E-G), again with no sex differences observed (males: 10.7 ± 3.1 days; females: 12.5 ± 4.6; t(16) = −0.288; *p* = 0.777). Performance in the full task over the five days before drug treatment is shown in H-K. No sex difference was found in the average performance on reward trials (males (N=7): 49.2±0.4 trials; females (N=11): 47.5±0.7 trials; t(16)= 1.733; *p* = 0.1). In the avoidance trials, on average, males successfully avoided the shock in more trials than did females (males (N=7): 47±1 trials; females (N=11): 31 ±6 trials; t(10.9) = 2.524; *p* = 0.03 (equal variances not assumed)).

#### Drug Administration in AAT

Seven male and 11 female animals reached criterion on both the cued-approach and cued-avoidance tasks and were put into the full task. The effect of FG7142 on performance in the full task was tested in these animals. FG7142 (Sigma-Aldrich, St. Louis, MO) was dissolved in sterile 0.09% saline with 15% v/v Tween 80 (Sigma-Aldrich, St. Louis, MO) for intraperitoneal injection of 1 mg/kg (FG1), 3 mg/kg (FG3), and 10 mg/kg (FG10). The vehicle for the control treatment was sterile 0.09% saline with 15% v/v Tween 80. Each animal received each of the four doses in a Latin square design over a period of seven days, with wash days (no injection) between each injection day. FG1, FG3, FG10, or vehicle was delivered 10 minutes prior to the beginning of the behavioral session. Animals were monitored for evidence of immobility or seizure.

#### Elevated Plus Maze after AAT

Anxiety-like behavior was quantified using the EPM (Harvard Apparatus 76-0074). Each animal was placed on the maze with their head facing an open arm. Video was captured with a FLIR Vue Pro thermal camera and analyzed with PanLab’s SMART video-tracking system. The time spent in the open and closed arms was measured and compared across groups. EPM was conducted in two separate cohorts of rats. The cohort of rats used in the AAT was tested on the EPM following the completion of operant testing, when the animals were free-feeding. A separate, naïve cohort was tested at age P73, also when the animals were free-feeding.

### Task 3: Cognitive Flexibility

#### Animals

Eight male and 8 female Long-Evans rats were delivered to the Oregon Health and Science University animal facility at the age of 60 days. Animals were housed with *ad libitum* food and water in temperature-controlled and humidity-controlled rooms on a 12 h on/12 h off reverse light/dark cycle with lights on at 1900 hours. After 1 week of acclimatization, animals were handled and weighed daily. Animals were food restricted to 80% of their free-feeding body weight. Animals were fed once daily after completion of behavioral tasks.

#### Training

For initial training, operant chambers were set up with a pellet dispenser and food trough on one wall and a nose poke (with light) on the opposite wall. On day 1, rats were habituated to the chambers. Approximately 20 sugar pellets were placed in the food trough and the nose poke port, and rats were allowed to freely explore the chamber for 30 mins. On days 2 and 3, rats began nose poke training. A nose poke was continuously available on the wall opposite the pellet dispenser, and each nose poke made by the rat delivered pellets on an FR1 schedule. On day 4, rats began training for the requirement to respond at the nose poke in less than 10 seconds. During this training, the port opened every 20 seconds, at the beginning of each trial. If rats responded within 10 seconds, a pellet was delivered and the nose poke port closed. If the rat did not respond within 10 seconds, no pellet was delivered and the nose poke closed until the beginning of the next trial. By day 8, rats were consistently responding at the nose poke port in less than 10 seconds and so moved on to side bias testing. On the final day of training, a side bias test was administered (modified from (Floresco, Block, & Tse, 2008) (Wallin & Wood, 2015)) Two nose poke ports were now available on the wall opposite the food trough. On each trial the rat could respond in either port to receive 1 sugar pellet. On subsequent trials, the rat was only rewarded for responding in the opposite port, and trials continued until the rat responded in both ports. This cycle was repeated 7 times. The nose poke port selected first on ≥4 of 7 cycles was considered the side bias. Nose poke port lights were not illuminated at any point during training procedures.

#### Testing

Set-shifting and reversal learning task procedures were modified from (Floresco et al., 2008). Rats were tested in daily sessions of 60 trials (20 seconds/trial). Each trial began with illumination of the house light. Three seconds later, one of the nose poke port lights was illuminated. A response made during this 3-second period between trial initiation and nose poke illumination was recorded and counted as a premature response, but was not punished. In a pair of trials, the left and right port lights were each illuminated once in random order. A correct response resulted in immediate delivery of 1 pellet, illumination of the food trough light, and retraction of the levers. The house light and trough light remained illuminated for 4 seconds while the pellet was retrieved. With an incorrect response or trial omission, the chamber immediately reverted to ITI. Choice latency was defined as the amount of time elapsed between the nose poke illumination and the rat’s response.

For initial discrimination learning, rats were assigned to the nose-poke port opposite their side bias (either left or right). Discrimination learning required rats to ignore the stimulus light and respond on their assigned side. Once criterion was met, the discrimination rule was reversed, such that rats were required to respond at the port on the opposite side. After completing the reversal, rats performed an extra-dimensional set-shift. This rule required rats to attend to the previously-irrelevant stimulus light, and to always choose the nose-poke port with the illuminated light. Criteria for success on each task (initial discrimination, reversal, and set-shift) was 8 consecutive correct responses with a minimum of 30 trials completed. If criterion was not met on the first day, rats continued to perform the same task again in daily sessions of 60 trials. Once criterion was met, rats moved on to the next task on the following day.

### Statistical Analyses

All data are expressed as mean ± standard error of the mean. The statistical analysis of behavioral data was performed using Student’s t-test when two groups were being compared. Task one results were analyzed using repeated measures ANOVA with sex as the between subjects factor and block as the repeated measure. Drug-behavior interaction was analyzed using two-way repeated measures analysis of variance (ANOVA). For each task in the cognitive flexibility experiment (initial discrimination, reversal learning and set-shifting), the number of trials required to reach criterion by male and female groups was compared by 2×2 mixed factorial ANOVA, with sex as the between subjects factor and task as the repeated measure. Likewise, premature responses and choice latency were averaged for each group in each task and compared by 2×2 mixed factorial ANOVA. Values of *p* < 0.05 were considered statistically significant, and values of 0.05 < *p* < 0.1 were considered trending.

### Open Practices Statement

The data and materials for all experiments will be made available through Open Science Framework upon acceptance. These experiments were not preregistered.

## RESULTS

### Sex differences during the Punishment Risk Task (PRT)

Goal directed behavior often requires balancing reward motivation with punishment avoidance. We modelled this by adding a component of probabilistic punishment to a reward-seeking task (Figure 1A), such that reward delivery was paired with a varying chance of foot shock.

Females exhibited a greater increase in response latency than males (Figure 1B). This suggests females are more affected by risk of punishment during reward seeking behavior. Combined across groups, rats displayed a significant increase in latency to nose-pokes as the probability of shock increased. There were no significant sex-or block-associated differences found in reward retrieval response times, suggesting similar levels of motivation for reward (Figure 1C).

### Sex differences in learning and performance on the AAT

Males and females acquired cued action for reward learning at similar rates (Figure 2B-D). In contrast, females acquired initial instrumental avoidance learning significantly faster than males, with females completing the hand-shaping of avoidance behavior in fewer sessions than males (males (N=7): 4.29±0.606 sessions, females (N=11): 2.55±0.207 sessions; t(7.425) = 2.717; p = 0.028). However, after hand-shaping, there was no sex difference in acquisition of cued avoidance learning between males and females (Figure 2E-G). After training was completed, rats were tested on the full AAT until behavior stabilized (defined as 5 consecutive days with no statistical difference between sessions). Behavior from these 5 days was averaged and shown in Figure 2H-K. Males and females did not differ in performance of approach trials (Figure 2H-I). In contrast, during avoidance trials males successfully avoided the shock more often than females (Figure 2J-K). This was surprising, as females had acquired the initial instrumental avoidance behavior (by hand-shaping) more quickly than males.

Upon closer examination, we found that avoidance behavior was not consistent across females but rather distributed in a bimodal fashion. A subset of the female animals exhibited an interesting behavior of approaching the correct instrumental action during the avoidance cue, but waiting until the shock began before immediately escaping the shock. This behavior suggested that these animals retained the learning of the cue-outcome and action-outcome relationships, but were not as motivated to completely avoid the shock. We found that the females could be divided into two subsets: ‘avoiders’ who responded during the avoidance cue in order to avoid receiving the shock entirely, and ‘escapers’ who waited through the avoidance cue until the shock began before responding to escape the shock. Avoiders (N=6) were categorized as those animals who performed above the 70% successful avoidance criterion and escapers (N=5) were those who performed below criterion. Female avoiders and escapers did not differ on any measure other than correct avoidance trials (t(9) = 6.51; *p* < 0.001), including correct rewarded trials (t(9) = 1.272; *p* = 0.2). These two female groups had also shown no difference in the number of hand-shaping days required to acquire the avoidance learning (avoiders: 2.5±0.3 days, escapers: 2.6±0.3 days; t(9) = −0.238, *p* = 0.818).

### Sex differences in different models of anxiety

Given that sex differences in the above tasks were more pronounced under the context of punishment or aversive variables, we conducted two experiments to gauge the impact of traditional models of anxiety in the context of our tasks. These included measuring (1) the dose-response effect of a pharmacological model of anxiety (anxiogenic drug FG7142) on performance of the AAT and (2) EPM performance of animals that had undergone training on the AAT compared to naïve animals.

Under FG7142 treatment, approach behavior diminished similarly in both sexes as dose increased, as indicated by fewer successfully completed approach trials (Figure 3A). FG7142 also increased latency to respond to the reward cue in both males and females (Figure 3B). However, latency to retrieve the sugar pellet reward was unaffected by FG7142 treatment, suggesting no effect on motivation to consume reward (Figure 3C).

**Figure 3:**
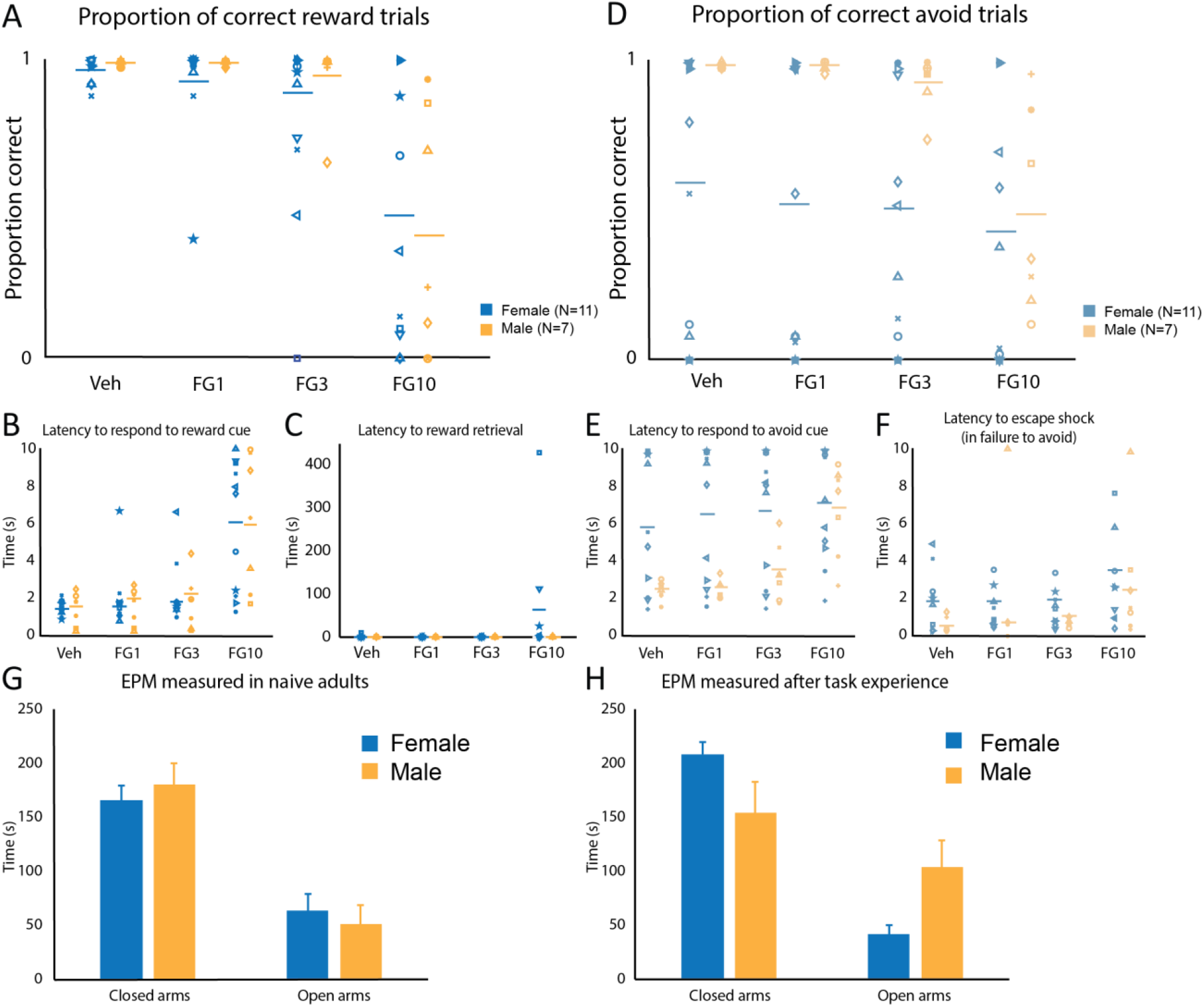
Sex differences in the Approach Avoid Task with administration of varying doses of FG7142. Anxiety induced by FG-7142 affected male and female performance equally on reward trials. There was a significant effect of dose on correct reward trials completed (RM-ANOVA; F(3, 48) = 17.465; *p* < 0.001), no significant effect of sex (F(1,16) = 0.373; *p* = 0.55), and no interaction between dose and sex (F(3, 48) = 0.559; *p* = 0.643) (A). A significant main effect of dose was found in the latency to respond to the approach cue (F(1,16) = 30.818; *p* < 0.001), but no main effect of sex (F(1, 16) = 0.096; *p* = 0.761) and no interaction between dose and sex (F(3, 48) = 0.101; *p* = 0.959) (B). No significant main effects of dose (F(3, 36) = 0.994; *p* = 0.4), sex (F(1, 12) = 1.431; *p* = 0.3), or interaction (F(3, 36) = 0.961; *p* = 0.4) were found in the latency to retrieve the sugar pellet reward (C). Avoidance behavior was diminished with administration of FG7142, as quantified by successfully avoided shock trials. RM-ANOVA indicated a significant effect of dose (F(3, 48) = 12.294; *p* < 0.001), a marginally significant effect of sex, with males showing greater sensitivity to FG7142 (F(1,16) = 4.269; *p* = 0.055), and a significant interaction between dose and sex (F(3, 48) = 4.782; *p* = 0.005) (D). Latency to respond to the avoidance cue was affected by dose of FG-7142 (RM-ANOVA; F(3, 48) = 12.643; *p* <0.001) and sex (F(1, 16) = 4.794; *p* = 0.04), and there was a significant dose * sex interaction (F(3, 48) = 5.107; *p* = 0.004) (E). In the cases where the animal did not avoid the shock, latency to escape the shock (by performing the avoidance action during the shock) is shown in panel F. Generalized anxiety-like behavior was assayed using the EPM. When tested as naïve adults, males and females showed similar behavior on the EPM. No difference was found between males and females in time spent in the closed arms (males: 181 ± 20 s (N=6); females: 165 ± 17 s (N=6); t(10) = 0.578; *p* = 0.576) or the open arms (males: 52 ± 13 s; females: 64 ± 16 s; t(10) = −0.567; *p* = 0.583) (G). After experience on the AAT, females spent more time than males in the closed arms (males: 154 ± 29 s (N = 7); females: 208 ± 12 s (N = 11); t(16) = −2.005; *p* = 0.062) and significantly less time than males in the open arms (males: 103 ± 26 s; females: 43 ± 8 s; t(16) = 2.609; *p* = 0.019) (H).

Although FG7142 treatment affected approach behavior similarly in males and females, it did elicit sex differences in avoidance behavior, measured as the proportion of avoidance trials in which the rat successfully avoided punishment. FG7142 significantly decreased the proportion of successful avoidance trials. There was a significant interaction between dose and sex, with FG7142 affecting male behavior to a greater extent than female behavior (Figure 3E). However, there was no effect of drug or sex on latency to escape shock during unsuccessful trials, suggesting that differences cannot be attributed to general changes in motor activity.

Anxiety-like behavior in the EPM was compared between males and females in a naïve cohort (Figure 3G) and in the cohort of animals used in the AAT after they had completed all behavior and drug administration testing and were free-feeding (Figure 3H). Among the animals that experienced the approach and avoidance behavior learning, females spent significantly more time in the closed arms and less time in the open arms than males. Female avoiders and escapers did not differ in time spent in either closed arms (avoiders: 211±10 seconds, escapers: 205±25 seconds; t(5.284) = 0.230, *p* = 0.826) or open arms (avoiders: 34±3, escapers: 53±17; t(4.192) = −1.089, *p* = 0.335). However, in the cohort of naïve animals tested on the EPM, there were no sex differences in in time spent in the closed arms or the open arms, suggesting an interaction between sex and experience of behavioral training and/or drug treatment on expression of generalized anxiety-like behavior.

### No sex differences in a reward-based cognitive flexibility task despite increased premature responding in males

The above data indicated that there is no sex difference in learning or maintenance of reward-guided behavior in the absence of punishment contingency. To determine if this lack of sex difference in rewarded behavior extends to tasks with a cognitive load, we compared male and female learning and performance of a cognitive flexibility task (Figure 4A). We found no sex differences in initial discrimination learning, reversal learning, and set-shifting (Figure 4B). There was no effect of sex on the number of trials required to reach criterion, but a significant effect of task on trials to criterion, with all rats requiring more trials on the reversal learning task than either the initial discrimination or the set-shift. There was no sex x task interaction, such that both males and females exhibited a similar pattern of performance to criterion over the three tasks. There were also no effects of sex or task on choice latency and no significant interactions on average latency to respond. The only sex difference we observed was in the number of premature nose poke responses (Figure 4C) with males making significantly more premature responses than females. However, there was no effect of task on this measure and no sex x task interaction.

**Figure 4:**
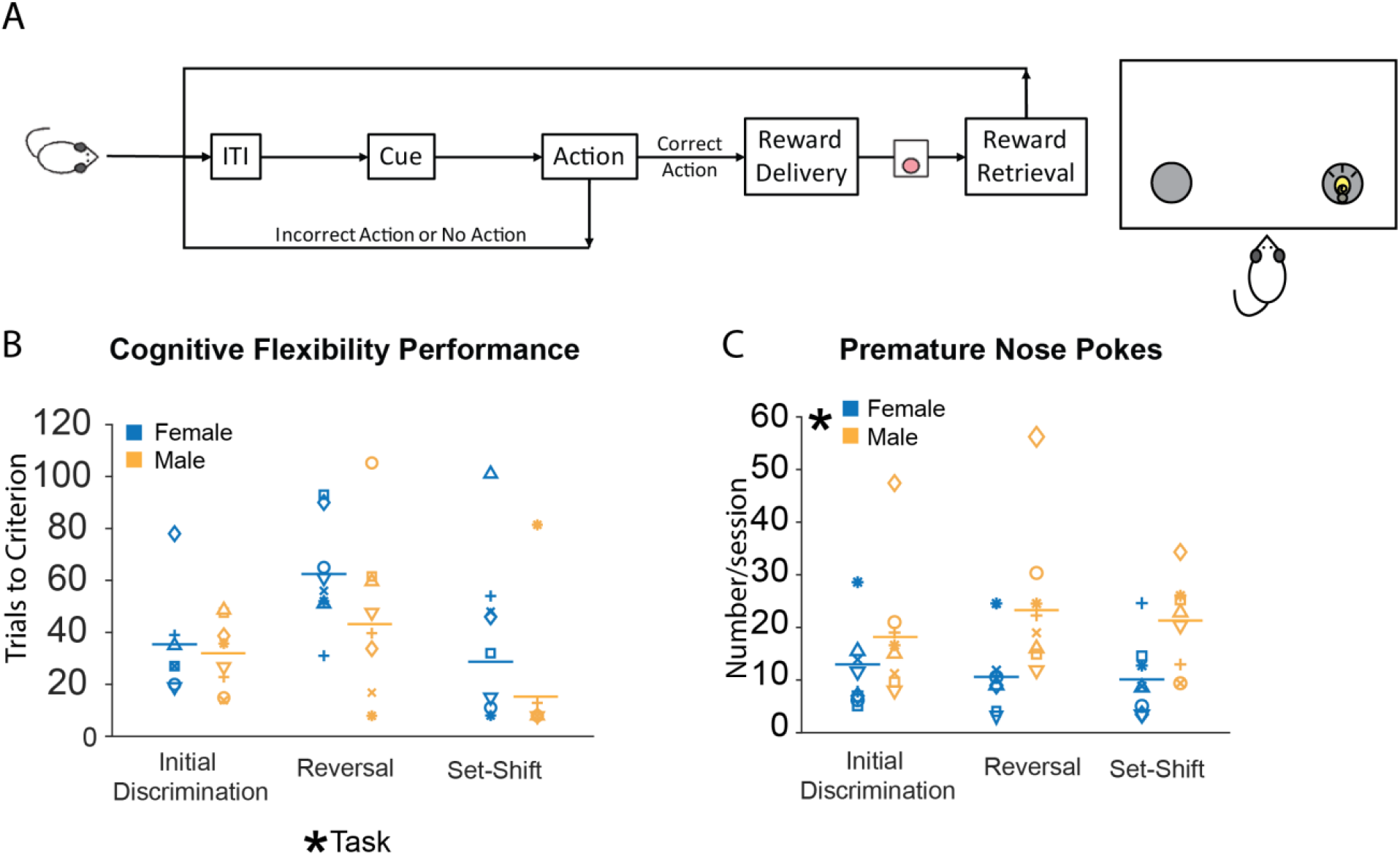
Sex differences in performance and premature responses during cognitive flexibility tasks (A). There was no effect of sex on performance (F1,14=2.44, p>0.05). However, there was a significant effect of task on performance, with all rats requiring more trials to reach criterion on the reversal learning task than either the initial discrimination or the set-shift (F2,28=7.42, p<0.05) (B). There was a sex difference in premature nose pokes, with males making significantly more premature responses than females (F1,14=5.45, p<0.05) (C). However, there was no effect of task on this measure and no sex x task interaction.

## DISCUSSION

Recent efforts to redefine diagnostic criteria for mental disorders have suggested an approach that explores the basic dimensions on which behavior can be described, cardinal of which are reward processing and punishment avoidance. Here we explored sex differences along these dimensions using multiple and distinct behavioral paradigms. We found a lack of sex differences in learning and execution of an appetitive instrumental action but faster punishment-avoidance learning in females. After learning, sex differences in aversive contexts depended on whether the risk of punishment was certain or unpredictable: females’ actions were more sensitive to punishment if it was presented with uncertainty (i.e. risk) but less sensitive to punishment if it could be avoided with certainty. This work provides novel behavioral assays to investigate brain mechanisms that account for nuanced sex differences in reward-seeking and punishment-avoidance behaviors.

### The study of sex differences relevant to psychiatric disease requires novel models

The current study developed novel models and employed a multifaceted investigation of motivated behavior in males and females. We assessed reward-motivated actions under different contexts, incorporating punishment, anxiogenic manipulations, or increased cognitive load. We detected no sex differences in acquisition or expression of purely reward-motivated actions, regardless of cognitive load. Sex differences, however emerged when the threat of punishment was introduced.

In the PRT, where reward seeking actions were accompanied with a risk of punishment, the response latency of both sexes increased, consistent with the expectation that threat-causes hesitation in reward-seeking actions. Females, however, exhibited a significantly greater increase in response latency with increasing risk of punishment than males. This suggests a higher sensitivity to the uncertain threat in females that is associated with the action. This is consistent with other observations indicating that female rats are more likely than males to avoid rewards associated with threatening stimuli. Females demonstrate reduced preference for risk-associated rewards during a risky decision-making task (Orsini et al., 2016), as well as increased latency to perform a risky choice. Female rats also display reduced foraging for food and water in a randomly punished context, despite both sexes demonstrating a comparable reduction in time spent in this context (Pellman, Schuessler, Tellakat, & Kim, 2017). More importantly, this mirrors observations of reward motivated behavior in humans, and suggests that increased responsiveness to risk of punishment (rather than blunted reward sensitivity) may be responsible for increased vulnerability to anxiety symptoms in females (Sheynin et al., 2014). Along the same lines, reduced responsiveness in risk of punishment in males is consistent with higher incidence of impulsivity in men (Perry & Carroll, 2008).

To further parse sex-differences in reward and aversion, we designed the AAT which assessed reward seeking and punishment avoidance in the same behavioral series. Again, we found sex differences only in avoidance behavior, with female rats acquiring avoidance behavior more rapidly than males, consistent with previous data showing that females learn more effectively in a shuttle box avoidance task (Dalla & Shors, 2009). However, once performance stabilized, males performed more successful avoidance trials than females, suggesting increased motivation to avoid punishing outcomes in males. Thus, sex differentially mediated acquisition and expression stages of avoidance behavior. Of note, this is the opposite of our observation in the PRT, in which punishment was uncertain and inescapable, suggesting that the anxiety-like response is dependent on the predictability, avoidability, and likelihood of the aversive outcome. These sex differences and individual differences in the context of avoidance suggest that, in general, response to punishment is less uniform than response to reward.

This sex difference in expression was partially influenced by a subset of females: while about half of females successfully avoided the shock ("avoiders”), the other half consistently waited for shock to begin before immediately performing the response to turn it off ("escapers”). This presence of an avoidance strategy unique to females has also been observed in a traditional fear conditioning task. Whereas male subjects show a near-uniform elevation in freezing to an aversive conditioned stimulus, a subset of females perform an active escape-like motor response termed “darting” (Gruene et al., 2015). Both this and the current study describe sex differences with a bimodal response profile within the female population. Here, we reveal that (1) these effects differ between Pavlovian fear conditioning and active avoidance tasks, and (2) this effect is valence specific, with sex differences and divergent female groups manifesting during punishment avoidance but not reward-seeking within the same behavioral context. Although it is difficult to determine why females appear to have evolved more varied and context-dependent avoidance behavior, this highlights the importance of further investigation into sex differences in these behaviors. In particular, these findings are highly relevant to pathological anxiety in humans, and suggest that understanding and treatment of this disorder will not be “one-size-fits-all” or even “one-size fits all males or all females.” Future studies will seek to elaborate on the neurobiological processes underlying these distinctions in avoidance.

In the context of the AAT, we also measured behavior on the EPM, which we interpret as a measure of generalized anxiety. We found that naïve rats showed no sex difference in behavior on the EPM, but after the experience of the AAT and FG7142 administration, females showed greater anxiety-like behavior on the EPM. Our results from the AAT suggest two related hypotheses: (1) females may be more resilient than males to an acute aversive event, and (2) continued experience of the aversive context may lead to increased anxiety in females, but not males. The first of these is consistent with the results of a study by Martin, et al. that showed that re-exposure to a context associated with pain results in pain hypersensitivity in male mice, but not female mice, and similarly in humans, that men, but not women, self-reported high levels of stress in a context previously associated with tonic pain (L. J. Martin et al., 2019). Considering these findings in interpreting our current results, it is possible that males experienced more aversive motivation after re-exposure in our AAT, and this could explain why (1) a subset of males exhibited learned helplessness-like behavior, and (2) the males in the full task were more motivated to avoid the shock on all avoidance trials. The second hypothesis is consistent with the findings of Day et al., who showed that females demonstrate greater discrimination than males after limited training, and generalization after extended training on a fear discrimination task (Day, Reed, & Stevenson, 2016). Together with our current findings, these results are consistent with the idea that women’s vulnerability to anxiety disorders stems from accumulating experience of aversive contexts

Finally, we investigated potential sex differences in reward-motivated actions within a context of high cognitive load, requiring flexible shifts between dynamic cue-action-reward contingencies. We found no sex differences in reward-motivated discrimination learning, nor in cognitive flexibility. While there was no difference in task performance, males made significantly more premature responses than females. Interestingly, behavioral disinhibition is a characteristic of several psychiatric diseases that preferentially affect males, such as ADHD, OCD, and substance abuse disorders. Therefore, it is possible that sex-specific differences in the neural circuitry supporting behavioral inhibition predispose males to these types of disorders (Perry & Carroll, 2008; Weafer, De Arcangelis, & de Wit, 2015; Weafer & de Wit, 2014). As reviewed in Weafer & de Wit, 2014, male laboratory animals consistently exhibit increased impulsive action relative to female subjects. Additionally, in humans, impulsive action is implicated in an increased risk for drug abuse and addiction (Perry & Carroll, 2008; Weafer et al., 2015).

Therefore, investigating sex differences in the neural basis of impulsivity will inform our understanding of sex differences in substance abuse and other psychiatric disorders.

### Evolutionary relevance of context-specific sex differences in motivated behavior

It is interesting to consider potential evolutionary relevance of context-specific sex-differences in motivated behavior. First, uniform behavioral responses suggest an optimal behavioral phenotype: When a reward is certain and involves no threat, maximal reward responding is advantageous. On the other hand, when reward is tempered by potential punishment there is no single optimal behavior; rather, there is a spectrum of phenotypes ranging from maximal reward response in spite of potential punishment, to the complete cessation of reward response due to punishment. Neither of these extremes are particularly useful: lack of punishment responsivity is likely to cause maladaptive risk taking whereas maximal punishment sensitivity will impede reward-motivated actions (and thus survival). When there is no single optimal strategy, it is advantageous for subgroups of behavioral phenotypes to emerge, equipping a species to survive in dynamic environments.

Our results, in combination with other literature (Orsini et al., 2016), suggest that there are sex differences in behavioral strategies to punishment risk and avoidance behavior. In particular, we found that avoidance motivation in females is context-dependent, with females more vulnerable to unpredictable punishment or under anxiety during new learning, and less vulnerable when punishment contingencies are well-learned and predictable. We also found that within a group of females, different strategies emerged in response to threat. In particular, the “escaper” female behavior of waiting for a shock to happen before immediately turning it off presents a unique opportunity for information gathering. Before escaping the shock they confirm that (1) the shock is still an existing threat and (2) the known method for escape still works. This may facilitate behavioral adaptation in females Perhaps females inhabit a more variable umwelt (with shifting priorities of self-survival and maternal behavior), and thus, it is more important to develop context-specific and adaptable avoidance responses. This continued information gathering by females is essentially prolonged exploratory behavior, which requires increased energy expenditure. Such an investment may not be beneficial enough to male survival to warrant the cost, and thus the behavior is not exhibited by males.

### Rethinking traditional models of anxiety

Emerging evidence demonstrates the importance of careful experimental design for the study of sex differences in behavior. For instance, when males and females were first compared in the Morris Water Maze (MWM; a standard test of spatial memory), females exhibited impaired spatial memory (measured as time taken to reach a hidden platform). However, subsequent studies noted that females exhibit thigmotaxis (an anxiety-related behavior), swimming along the edge of the maze rather than straight across it (thus taking longer to reach the platform). Habituation of both sexes to the maze set-up decreases thigmotaxis in females, and eliminates sex differences in spatial memory (Perrot-Sinal, Kostenuik, Ossenkopp, & Kavaliers, 1996; Roof & Stein, 1999). Likewise, in the current study we found that anxiety affects behavior differently in males and females, depending on the specific parameters of the task.

Design considerations are also apparent for the EPM, a standard assessment of anxiety in rodents (Hogg, 1996; Pellow, Chopin, File, & Briley, 1985). We initially found no sex difference in anxiety with the EPM, despite well-known sex differences in anxiety-related psychiatric illness. Thus, we should not conclude that there are no sex differences in anxiety, nor that the EPM is useless, but rather that the EPM is an invalid measure of the anxiety processes relevant to sex differences in mental illness. This finding is in line with previous criticisms of the EPM. For instance, some antidepressants that are clinically effective for reducing anxiety fail to exert anxiolytic effects in the EPM. Additionally, handling history has been shown to effect EPM results (Andrews & File, 1993). Likewise, in the current study we found increased anxiety in females only after a history stressful behavioral testing. Other studies have shown similar sex-dependent responses to stress. For instance, prenatal stress affects pups differently depending on sex, with males and females exhibiting different anxiety behaviors as adults (Schulz et al., 2014), as well as sex-dependent effects on the underlying neurobiology (Schulz et al., 2013). Studies in male rodents have shown differential effects of escapable and inescapable stress in the amygdala (Weinberg et al., 2010) and hippocampus (Shors, Seib, Levine, & Thompson, 1989), both of which are brain regions shown to have sex-specific effects of stress (Cahill et al., 2001; Shors, Chua, & Falduto, 2001). As with the MWM, these results caution against generalizing the results of one task to the conclusions of sex differences in a certain modality. Rather, investigation of relevant sex differences in behavior requires careful model design and interpretation.

## CONCLUSION

We characterized sex differences in constructs of cognition and motivated behavior related to psychiatric disorders. We observed sex differences in reward seeking actions that were specific to the contexts of punishment or anxiety. Other task variables, such as cognitive load, turned out to be irrelevant domains to reveal sex differences. Future studies can take advantage of these tasks to probe the mechanisms underlying sex differences in emotional processing and related disorders.

